# Impact of different intensities of intermittent theta burst stimulation on the cortical properties during TMS-EEG and working memory performance

**DOI:** 10.1101/157917

**Authors:** Sung Wook Chung, Nigel C. Rogash, Kate E. Hoy, Caley M. Sullivan, Robin F. H. Cash, Paul B. Ftizgerald

## Abstract

**Introduction:** Intermittent theta burst stimulation (iTBS) is a non-invasive brain stimulation technique capable of increasing cortical excitability beyond the stimulation period. Due to the rapid induction of modulatory effects compared to conventional repetitive transcranial magnetic stimulation (rTMS) paradigms, prefrontal application of iTBS is gaining popularity as a therapeutic tool for psychiatric disorders such as depression. In an attempt to increase efficacy, higher than conventional intensities are currently being applied. The assumption that this increases neuromodulatory effect is well established for the standard rTMS paradigms but may be mechanistically false for iTBS. This study examined the influence of intensity on the neurophysiological and behavioural effects of iTBS in the prefrontal cortex.

**Methods:** 16 healthy participants received iTBS over prefrontal cortex (F1 electrode) at either 50, 75 or 100% resting motor threshold (rMT) in separate sessions. Single-pulse TMS and concurrent electroencephalography (EEG) was used to assess changes in cortical reactivity measured as TMS-evoked potentials (TEPs) and TMS-evoked oscillations. The n-back task (2-back and 3-back) were used to assess changes in working memory (accuracy and reaction time).

**Results:** The data can be summarised as an inverse U-shape relationship between intensity and iTBS plastic effects, where 75% iTBS resulted in the largest neurophysiological changes both in TMS-EEG and working memory. Even though 75% iTBS showed significantly decreased reaction time in the 3-back task, between condition comparisons revealed no significant differences.

**Conclusions:** The assumption that higher intensity results in greater neuromodulatory effects is false, at least in healthy individuals, and should be carefully considered for clinical populations. Neurophysiological changes associated with working memory following iTBS suggest functional relevance. However, the effects of different intensities on behavioural performance remain elusive in the present healthy sample.

**Highlights:**

- Effects of prefrontal iTBS at 50, 75 and 100% rMT were investigated
- Inverse U-shape relationship between intensity and neurophysiological effects
- Effects on the behavioural performance remain elusive in healthy individuals

**Disclosures and conflict of interests:** SWC was supported by a Monash Graduate Scholarship. NCR is supported by a NHMRC Early Career Fellowship (1072057). KEH is supported by a NHMRC Career Development Fellowship (1082894). PBF is supported by a NHMRC Practitioner Fellowship (1078567). PBF has received equipment for research from MagVenture A/S, Medtronic Ltd, Cervel Neurotech and Brainsway Ltd and funding for research from Neuronetics and Cervel Neurotech. He is on the scientific advisory board for Bionomics Ltd. There are no other conflicts.

## 1. Introduction

Repetitive transcranial magnetic stimulation (rTMS) is a non-invasive brain stimulation technique capable of modulating cortical activity beyond the stimulation period. Clinical applications of rTMS have been studied in various neurological and psychiatric disorders (Machado et al., 2013), especially in the treatment of depression (George et al., 2010; George et al., 2013; O'Reardon et al., 2007). Recently, a modified form of rTMS known as theta-burst stimulation (TBS), has been investigated as a potential treatment of depression, with promising therapeutic effects (Chung et al., 2015a; Duprat et al., 2016; Li et al., 2014). TBS was modified from an animal stimulation paradigm (Larson et al., 1986) and elicits long-term potentiation or depression (LTP/LTD) – like changes depending on the stimulation pattern in human (Huang et al., 2005). In the motor cortex, intermittent TBS (iTBS, 2sec on, 8 sec off, 600 pulses, 3 minutes duration) elicits LTP-like increases in cortical excitability whereas continuous TBS (cTBS, 40 sec on, 600 pulses) evokes LTD-like decreases cortical excitability (Huang et al., 2005; Suppa et al., 2008). Due to the rapid induction of modulatory effects compared to conventional rTMS, TBS is an attractive option for neuromodulatory treatments in clinical disorders (Chung et al., 2015a; Machado et al., 2013). For psychiatric conditions, this typically involves stimulation delivered to prefrontal cortical regions such as dorsolateral prefrontal cortex (DLPFC) (Desmyter et al., 2016; Li et al., 2014; Plewnia et al., 2014). As adoption of TBS increases in the clinical literature, the lack of consensus with respect to optimal intensity is becoming increasingly evident. Conventionally, TBS in the motor cortex has been applied at 80% of active motor threshold (aMT) (Huang et al., 2005), equivalent to approximately 75% of resting motor threshold (rMT). Recent reports of the stimulation intensity used in prefrontal TBS for therapeutic intervention have varied quite substantially in the range of 80 – 120 % of rMT (Bakker et al., 2015; Desmyter et al., 2016; Duprat et al., 2016; Li et al., 2014; Plewnia et al., 2014; Prasser et al., 2015). The underlying assumption is that the efficacy of iTBS will be greater with increasing intensity of stimulation. This is partially supported by linear responses to increases in the intensity of conventional rTMS in healthy individuals (1 Hz (Nahas et al., 2001)) and in clinical populations (10 Hz (Padberg et al., 2002)). Studies using different modulatory paradigms have also shown a shift from LTD- to LTP-like effects at higher intensity (Batsikadze et al., 2013; Cash et al., 2017a; Doeltgen and Ridding, 2011), corroborating the idea of increased propensity for LTP-like changes at higher intensity. However, systematic investigation of intensity-dependent effects of iTBS in the prefrontal cortex has not been established.

Another key question concerns the use of TBS for cognitive disorders. TBS was originally developed to mimic the natural firing patterns of neurons in the hippocampus, where high-frequency gamma oscillations (30 – 80 Hz) were modulated by the phase of lower frequency theta oscillations (4-7 Hz) (Lisman and Jensen, 2013). Applying electrical stimulation to the hippocampus with gamma frequency bursts nested in theta frequency rhythms resulted in robust long-term potentiation (LTP) (Larson et al., 1986). A similar theta-gamma coupling relationship in endogenous brain activity has been observed in human studies using electroencephalography (EEG) during cognitive functions (Lisman, 2010). It is therefore of particular interest whether iTBS in human can facilitate cognitive and memory processes, and to what extent the plastic changes elicited by TBS translate to changes in neurophysiological metrics of cognition and behavioural performance outcomes.

Recent advances in technology have enabled the measurement of plastic neuronal changes following neuromodulatory paradigms using concurrent recording of electroencephalographic responses to TMS (TMS-EEG) (Chung et al., 2015b; Farzan et al., 2016; Hill et al., 2016). Each TMS pulse elicits a TMS-evoked EEG response, and the change in the amplitude of TMS-evoked potentials (TEPs) and the power of TMS-evoked oscillations following TBS provide metric of plasticity in the prefrontal cortex (Chung et al., 2017). TEPs are composed of several components which are thought to represent excitatory and inhibitory postsynaptic potentials. A negative trough at a latency of approximately 100 ms (N100) has been associated with inhibitory mechanisms in motor (Bonnard et al., 2009; Premoli et al., 2014; Rogasch et al., 2013a) and prefrontal cortex (Chung et al., 2017; Rogasch et al., 2015), and is considered to be the most robust TEP component with the best signal to noise ratio (SNR) (Noda et al., 2016). Modulation of this component has been observed following TBS over the prefrontal cortex (Chung et al., 2017) and cerebellum (Casula et al., 2016; Harrington and Hammond-Tooke, 2015). Recent studies also suggest that a peak at a latency of 60 ms (P60) may be a correlate of neuronal excitability in motor cortex (Cash et al., 2017b) and DLPFC (Hill et al., 2017).

In the present study, we examined the relationship between iTBS intensity (50, 75 and 100% of individual rMT), LTP-like neural plasticity and the relationship to neurophysiological and behavioural metrics of learning and memory using N-back task (Haatveit et al., 2010). We hypothesized that iTBS would be accompanied by plastic changes in N100 and P60 amplitude. Secondly, we anticipated that the efficacy of iTBS would increase with increasing intensity. Thirdly, we hypothesized that these changes will be mirrored by increasing working memory (WM) performance measured via N-back task and neurophysiological correlates. The modulation of theta and gamma oscillatory activity was of particular interest since these frequency bands are targeted by TBS and involved in WM.

## 2. Material and methods

### 2.1. Participants

Sixteen healthy volunteers (7 female, 27.8 ± 8.6 years of age, 16.25 ± 2.11 years of formal education) participated in the study. All subjects were right-handed according to the Edinburgh Handedness Inventory, and the mini international neuropsychiatric interview (MINI) was performed (Sheehan et al., 1998) to confirm no history of mental illness. No participants were smokers. All participants provided informed consent prior to the experiment and the experimental procedures were approved by the Alfred Hospital and Monash University Human Research Ethics Committees.

### 2.2. Procedure

Each participant attended 3 sessions receiving iTBS at either 50%, 75% or 100% of their resting motor threshold (rMT). Each session was at least 72 hours apart and the session order was pseudorandomized across participants. The experimental procedures comprised of recording EEG during 50 single TMS pulses before (BL – baseline), 5 min post (T5) and 30 min post (T30) iTBS (Fig 1A). The N-back WM task (2-back and 3-back conditions) was also performed pre (BL) and 15 min post (T15) iTBS with concurrent EEG recording.

**Figure 1.**
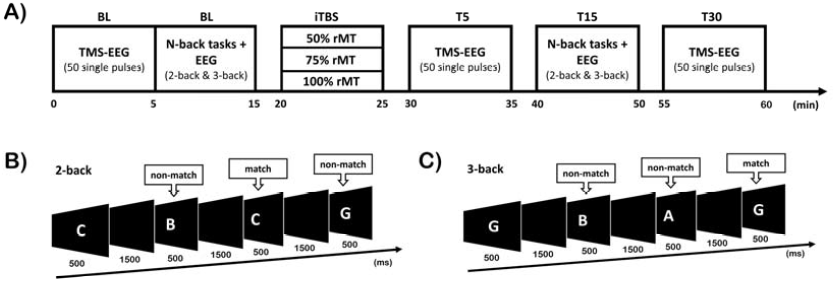
Schematic diagram of the experimental design. (A) Concurrent recording of electroencephalogram during transcranial magnetic stimulation (TMS-EEG) and N-back task were performed at baseline (BL). Intermittent theta burst stimulation (iTBS) was then administered at one of three intensities. TMS-EEG was rerepeated at T5 and T30 following iTBS, and the N-back at T15 following iTBS. (B-C) Diagrams illustrating trials of match and non-match during 2-back and 3-back tasks.

### 2.3. EEG recording

EEG was recorded with TMS-compatible Ag/AgCl electrodes and a DC-coupled amplifier (SynAmps2, EDIT Compumedics Neuroscan, Texas, USA). 42 electrodes were used on a 64-channel EEG cap (AF3, AF4, F7, F5, F3, F1, Fz, F2, F4, F6, F8, FC5, FC3, FC1, FCz, FC2, FC4, FC6, T7, C5, C3, C1, Cz, C2, C4, C6, T8, P7, P5, P3, P1, Pz, P2, P4, P6, P8, PO3, POz, PO4, O1, Oz, O2), and electrooculography recording was obtained with 4 electrodes, one positioned above and one below the left eye and one on lateral to the outer canthus of either eye. Electrodes were online referenced to CPz and grounded to FPz with exception to the lateral eye electrodes which were referenced to each other. For TMS-EEG, data were recorded with a high acquisition rate (10,000 Hz) using a large operating range (± 200 mV) to avoid amplifier saturation. Signals were amplified (1,000 x) and low pass filtered (DC-2,000 Hz). For EEG recording during N-back task, AC acquisition setting was used and the signals were filtered (low pass at 200 Hz, high pass at 0.05 Hz) and sampled at 1,000 Hz with an operating window of ± 950 μV. Electrode impedance levels were kept below 5 kΩ throughout the experiment. During TMS-EEG recording, subjects listened to white noise through intra-auricular earphones (Etymotic Research, ER3-14A, USA) to limit the influence of the auditory processing of the TMS click. The sound level was adjusted for each individual subject until single-pulse TMS at 120% rMT was barely audible.

### 2.4. Transcranial magnetic stimulation

Participants sat comfortably with their arms resting on a pillow throughout the experiment. The EEG cap was mounted following the 10-20 standard system and the resting motor threshold (rMT) was obtained from left motor cortex, which was identified as the minimum intensity required to evoke at least 3 out of 6 motor evoked potentials (MEPs) > 0.05 mV in amplitude (Conforto et al., 2004) via Ag/AgCl electromyography electrodes attached to the first dorsal interosseous (FDI) muscles. TMS was administered to the left prefrontal cortex at F1 electrode using 10/20 method of placement. The F1 electrode sits over the superior frontal gyrus with Brodmann area (BA) of 6, 8 and 9 (Koessler et al., 2009), and therefore, is part of dorsolateral prefrontal cortex. This electrode was chosen to minimize stimulation of scalp muscles which result in large artefacts in EEG recordings lasting up to 40 ms following the TMS pulse (Rogasch et al., 2013b). By minimising artefacts, the amount of correction needed in post-processing for the TMS-EEG signals is reduced. A MagVenture B-65 fluid-cooled coil (a figure-of-eight coil; MagVenture A/S, Denmark) was used for both single-pulse stimulation and iTBS (biphasic pulses, antero-posterior to postero-anterior current direction in the underlying cortex). The coil was positioned at 45° angle relative to midline, which has been shown to produce strongest stimulation in the prefrontal cortex (Thomson et al., 2013). A line was drawn on the coil at 45° angle, which would then sit perpendicular to the midline of the EEG cap to ensure same angle positioning. In addition, the edge of the coil was marked on the cap to reliably re-position the coil within 5 mm (Rogasch et al., 2013b).

Participants received 50 single pulses to left prefrontal cortex at an interval of 5 s (10% jitter) at 120% rMT before and after different stimulation intensities of iTBS. These parameters were chosen to be consistent with our previous study (Chung et al., 2017). Across different sessions, participants received iTBS at different intensities (50%, 75%, or 100% rMT). With the exception of intensity, iTBS parameters adhered to the originally described method (Huang et al., 2005). iTBS consisted of a burst of 3 pulses given at 50 Hz repeated at a frequency of 5 Hz, with 2 s of stimulation on and 8 s off repeated for a total of 600 pulses. The average stimulation intensity was as follows (% of maximum stimulator output; mean ± SD): 50% condition = 28.5 ± 3.0 %; 75% condition = 42.5 ± 5.3 %; 100% condition = 57.19 ± 5.6 %.

### 2.5. Working memory task

Participants were assessed on the N-back task with 5 mins of 2-back and 5 mins of 3-back conditions in a pseudorandomised order. Letters were in a random series of A to J, and participants were requested to respond with a button press when the presented letter was the same as the letter appeared either 2 trials (Fig 1B; 2-back) or 3 trials (Fig 1C; 3-back) earlier. Each letter was presented in white on a black screen for 500 ms with a 1500 ms interstimulus interval. Each N-back task consisted of 130 trials with 25% targets. Due to technical failure, data was not collected from one participant (complete data from 15 participants; 27.3 ± 8.7 years, 7 female). WM performance was assessed via accurate reaction time and d prime sensitivity index (d’) (z-transformed values of hit-minus false-alarm rates) (Haatveit et al., 2010).

### 2.6. EEG data preprocessing

EEG data were analysed offline using EEGLAB (Delorme and Makeig, 2004), FieldTrip (Oostenveld et al., 2011), TESA (Rogasch et al., 2016) and custom scripts on Matlab platform (R2015b, The MathWorks, USA). For TMS-EEG, data were epoched around the test TMS pulse (−1,000 to 1,000 ms), baseline corrected to the TMS-free data (−500 to −50 ms), and data around the large signal from TMS pulse (−5 to 10 ms) were removed and linearly interpolated. The epoched TMS-EEG data from all three time points (BL, T5, T30) were concatenated and analysed concurrently to avoid bias in component rejection. Data were downsampled to 1,000 Hz and visually inspected to remove epochs with excessive noise (i.e. muscle artefact), and bad channels (i.e. disconnected). An average of 47.6 (± 2.7) trials was included in the 50% iTBS condition, 47.4 (± 2.8) trials in the 75% iTBS condition and 48.0 (± 2.7) trials in the 100% iTBS condition across each time point. Two rounds of independent component analysis (FastICA algorithm using the ‘tanh’ contrast function) were applied to the data; the first to remove large amplitude muscle artefacts, and the second to remove other common artefacts following offline filtering. The first round of independent component analysis (ICA) used a semi-automated component classification algorithm (tesa_compselect function) to remove the remainder of the muscle artefact (Korhonen et al., 2011) (classified if component time course 8 times larger than the mean absolute amplitude across the entire time course). All data were bandpass filtered (second-order, zero-phase, Butterworth filter, 1-80 Hz) and bandstop filtered (48-52 Hz; to remove 50 Hz line noise) and epochs were inspected again to remove any anomalous activity in the EEG trace. The second round of FastICA was conducted, and additional artefactual components were removed based on a previous study (Rogasch et al., 2014) and using TESA toolbox as a guide (Rogasch et al., 2016). Components representing the following artefacts were removed; eye blinks and saccades (mean absolute z score of the two electrodes larger than 2.5), persistent muscle activity (high frequency power that is 60% of the total power), decay artefacts and other noise-related artefacts (one or more electrode has an absolute z score of at least 4).

For EEG during N-back tasks, data were epoched around the correctly encoded and maintained trials (−1,450 to 1,990 ms), and baseline corrected (−350 to −50 ms). Trials containing a button response in the epochs were excluded to avoid confounds introduced by motor preparation. Epoched EEG data for two time points (BL, T15) and two N-back tasks were concatenated and analysed concurrently to avoid bias in rejecting components. Data were visually inspected to remove epochs with excessive noise, and bad channels removed. An average of trials included in 50% iTBS conditions were – 76.6 (± 12.9) for 2-back, 74.5 (± 22.2) for 3-back; in 75% iTBS conditions were – 74.6 (± 13.3) for 2-back, 78.1 (± 17.8) for 3-back, and in 100% iTBS conditions were – 75.8 (± 14.3) for 2-back, 74.4 (± 24.5) for 3-back tasks. It has been demonstrated that late ERP components such as P300 encounters the risk of being distorted following high-pass filter above 1 Hz (Rousselet, 2012). However, drift in data filtered at 0.1 Hz is not suitable for ICA (Debener and De Vos, 2011). Therefore, steps were taken to minimize the distortion of ERPs; 1) All data were bandpass filtered (second-order, zero-phase, Butterworth filter, 0.1-80 Hz) and bandstop filtered (48-52 Hz), and set aside. 2) Original data were bandpass filtered at 1 – 80 Hz, and FastICA with artefact component removal was conducted as described above (only one round of ICA). 3) The ICA weight matrix from step 2 was then applied to the data in step 1.

For all EEG data, removed channels were interpolated, and data were re-referenced to common average reference. Finally, data were separated into time point blocks (TMS-EEG: BL, T5 and T30; N-back EEG: BL and T15), conditions (50%, 75% and 100% iTBS) and/or tasks (2-back and 3-back).

### 2.7. TMS-evoked potentials (TEPs) and event related potentials (ERPs) during N-back tasks

TEPs and ERPs were analysed using a global scalp analysis (cluster-based permutation statistics) to access the effect of iTBS across the cortex. For TEPs, the MagVenture stimulator has shown to introduce unwanted artefacts on electrodes in contact with the coil (Rogasch et al., 2013b). As such, the FCz electrode was chosen for TEP waveform representation. Amplitudes of TEPs were compared across time and conditions within pre-determined time window for N45 (30 – 55 ms), P60 (55 – 80 ms), N100 (90 – 140 ms) and P200 (160 – 240 ms). These peaks are known to occur following prefrontal TMS-EEG (Chung et al., 2017; Hill et al., 2017; Rogasch et al., 2015; Rogasch et al., 2014). A signal-to-noise ratio (SNR) analysis was performed on the average of three fronto-central electrodes (FC1, FCz, FC2) for each individual to validate the limited number of TMS pulses available for the analyses (∼ 47 pulses). The SNR was calculated by dividing the peak amplitude by the standard deviation (SD) of the TEPs in the pre-stimulus period (−500 to −50 ms) (Chung et al., 2017; Hu et al., 2010).

For ERPs during N-back tasks, the same electrode was used for graphical representation, and peaks were statistically compared within time window for N100 (70 – 110 ms), P150 (120 – 180 ms), N200 (190 – 260 ms) and P300 (280 – 380 ms) during encoding / maintenance period. These peaks were chosen for the implication of these peaks in visual WM tasks (Coull, 1998; Kok, 2001; Vogel and Luck, 2000).

### 2.8. TMS-evoked oscillations and event related oscillations during N-back tasks

TMS-evoked oscillatory power and event related oscillations during N-back tasks were measured by converting TEPs and ERPs into the frequency domain using Morlet wavelet decomposition (3.5 oscillation cycles (Casula et al., 2016; Hill et al., 2016; Hoy et al., 2015; Rogasch et al., 2015) with steps of 1 Hz between 2 Hz and 50 Hz, 10 ms time resolution on each trial for each electrode. The oscillatory power was then averaged to compute the total power of activity, which contained both evoked and induced oscillations. In line with recent discussions on the different approaches to the analysis of oscillatory activity in TMS-EEG (Pellicciari et al., 2017b), we explored the effects of iTBS on evoked neural oscillations alone. Normalised oscillatory power was then obtained by dividing all power bins by a mean baseline value (−650 to −350 ms). This baseline window was chosen to avoid the temporal smearing of post-stimulus activity into the baseline as the lowest frequency of interest (i.e. 5 Hz – 200 ms) would require at least 350 ms (3.5 oscillation cycles x 200 ms (5 Hz) = 700 ms; Half of the wavelet length – 700/2 = 350 ms). Power values were averaged in frequency bands of interest; theta (5 – 7 Hz) and gamma (30 – 45 Hz), and in time (50 – 250 ms for theta, 50 – 150 ms for gamma) prior to the computation of cluster-based statistics. Focused analyses were conducted on theta and gamma frequencies as theta-burst stimulation is comprised of these two frequency bands, and also due to the implication of synaptic plasticity by the interaction between theta and gamma frequency bands (Zheng and Zhang, 2015). For the N-back tasks, oscillations were investigated in two blocks; during the letter presentation (50 – 450 ms) and after the letter presentation (550 – 950 ms), and averaged across these time windows for both theta and gamma oscillations prior to the cluster-based statistics. Similar to the examination of oscillatory activity during TMS-EEG, evoked oscillations were also investigated.

Additional multi-dimensional cluster-based statistics were performed [time (50 – 500 ms for TMS-EEG; 50 – 950 ms for N-back tasks) x frequency (5 – 45 Hz) x space], as it is recommended to analyse the data in all possible dimensions (van Ede and Maris, 2016). Further subgroup analyses on alpha (8 – 12 Hz) and beta (13 – 29 Hz) bands were also conducted to explore any iTBS-induced change in these frequencies.

### 2.9. Source estimation

In order to establish the spread of activity following single-pulse TMS on F1 electrode, source estimation was performed. All source localisation was performed using depth-weighted minimum norm estimation (MNE) implemented in Brainstorm software (Tadel et al., 2011) which is documented and freely available for download online under the GNU general public licence (http://neuroimage.usc.edu/brainstorm/). A template anatomy (ICBM 152) in Brainstorm software was used as individual MRI scans were not obtained. The forward model used the Symmetric Boundary Element Method provided by OpenMEEG (Gramfort et al., 2010), and the inverse model was computed with dipole orientations constrained to be normal to the cortex.

### 2.10. Statistics

Statistical analysis was performed in SPSS (Version 22) and Matlab. Data did not meet the requirement for normality (Shapiro-Wilk test) in behaviour measures, and therefore nonparametric statistics were used. The Wilcoxon signed-rank tests were conducted for comparison between different pre- and post-iTBS measures to assess whether iTBS conditions altered WM performance. To assess whether iTBS conditions differentially affected working WM, Friedman’s Analysis of Variance by Ranks was used with a factor of condition (50%, 75%, 100%) to compare the change-from-baseline scores (post – pre; Δ) between conditions.

For analysis of electrophysiological data, non-parametric cluster-based statics were used (Oostenveld et al., 2011). Monte Carlo p-values were calculated on 5000 random permutations and a value of p < 0.05 was used as the cluster-statistical significance for all analyses, controlling for multiple comparisons across space and time (p < 0.025; two-tailed test). Within condition comparison was first conducted over time to assess whether iTBS conditions altered peak amplitudes / oscillatory power over time (post-iTBS vs pre-iTBS). To assess whether iTBS conditions differentially modulated these measure, Δ values (post – pre) were calculated and compared between conditions.

To assess the relationship between the changes in TMS-evoked activities, N-back related electrophysiology and WM performances, Spearman’s rank correlations were used.

## 3. Results

### 3.1. Single-Pulse TMS

An overview of TEP waveforms following single-pulse TMS over left prefrontal cortex (F1 electrode) and the source estimation at the peaks of interest (N45, P60, N100 and P200) are illustrated in Fig 2. The scalp topography and the source estimation of these peaks conform to other TMS-EEG studies in the prefrontal cortex (Chung et al., 2017; Hill et al., 2017; Rogasch et al., 2014).

**Figure 2.**
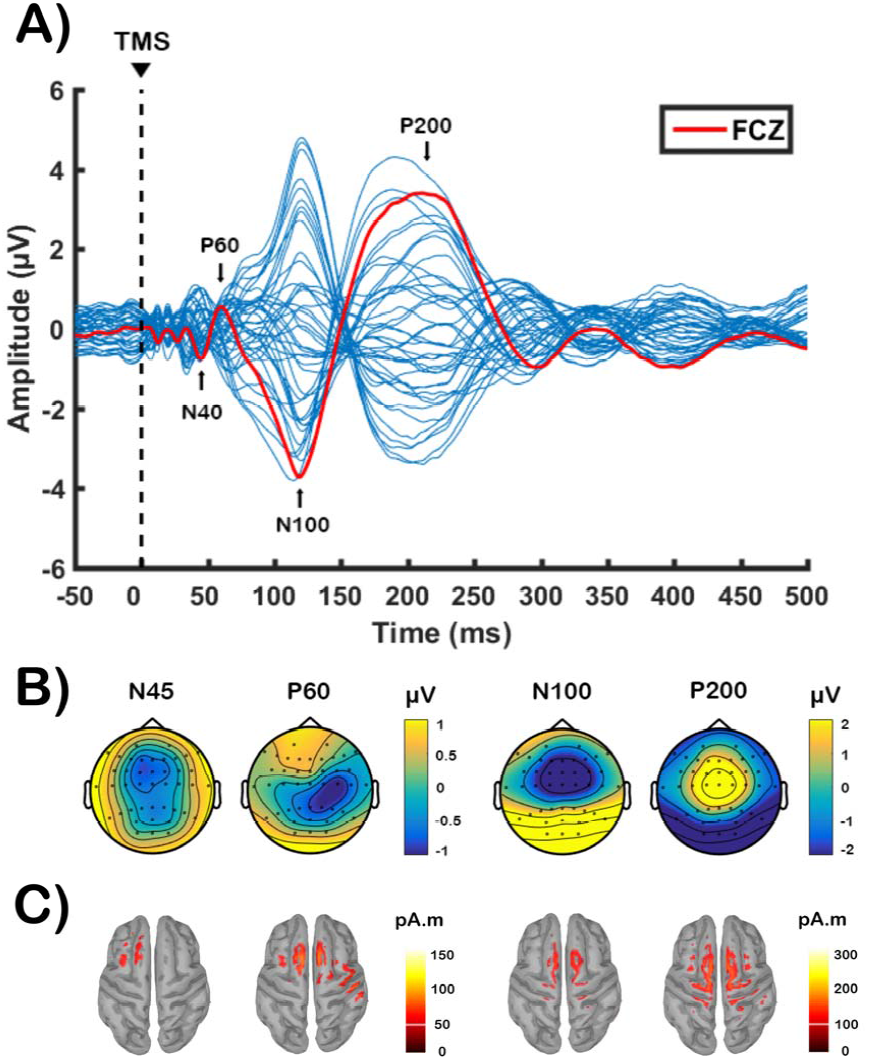
Transcranial magnetic stimulation (TMS)-evoked potentials following single-pulse stimulation over left prefrontal cortex (F1 electrode) before theta-burst stimulation (data combined across conditions at baseline). (A) Butterfly plot of all electrodes with peaks of interest (N45, P60, N100, P200) shown in text. The red line indicates the waveform obtained from FCz electrode for graphical representation. (B) Voltage distribution and (C) Minimum Norm Estimates (MNEs) of the source level activity at the cortex for each peak of interest.

The analysis of SNR can be found in the Supplementary Material, Table S1. Qualitatively N45 peaks showed moderate values (∼2.5 SDs), but other peaks, especially latter peaks (N100 and P200) showed high/acceptable SNR.

### 3.2. The effect of different iTBS intensities on TMS-evoked activity

We first assessed the after-effects of iTBS by comparing the amplitudes of TEPs over time, and across conditions. Using the cluster-based permutation tests between pre-iTBS (BL) and 5-min post iTBS (T5), we found that both 50% (0.115 – 0.140 ms, *p* = 0.011, right frontal; Fig 3A) and 75% iTBS (0.110 – 0.140 ms, *p* = 0.010, bilateral frontal; Fig 3B) resulted in an increased N100 amplitude. This change, however, was absent following 100% iTBS (*p* > 0.025; Fig 3C), and no other peaks showed any significant changes (all *p* > 0.025). We compared the TEPs between BL and 30-min post iTBS (T30), but no significant persistent effect remained (all *p* > 0.025). In order to evaluate the differences between conditions, iTBS-induced changes in TEP amplitude were calculated (post – pre) and compared. As our experimental design was not sham-controlled, this method of comparison would minimize the confounding factor (e.g. change over time unrelated to stimulation). We found that the change in N100 amplitude (Δ N100) was the largest with 75% iTBS, but less following 100% iTBS (75% > 100% iTBS: T5, 0.112 – 0.140 ms, *p* = 0.008), which was observed in fronto-central sensors (Fig 3D). Source estimation of N100 in these stimulation conditions supported the findings of the scalp-level analyses, where increased electrical activity was found in fronto-central region following 75% iTBS, but minimal change was seen following 100% iTBS (Fig 3D).

**Figure 3.**
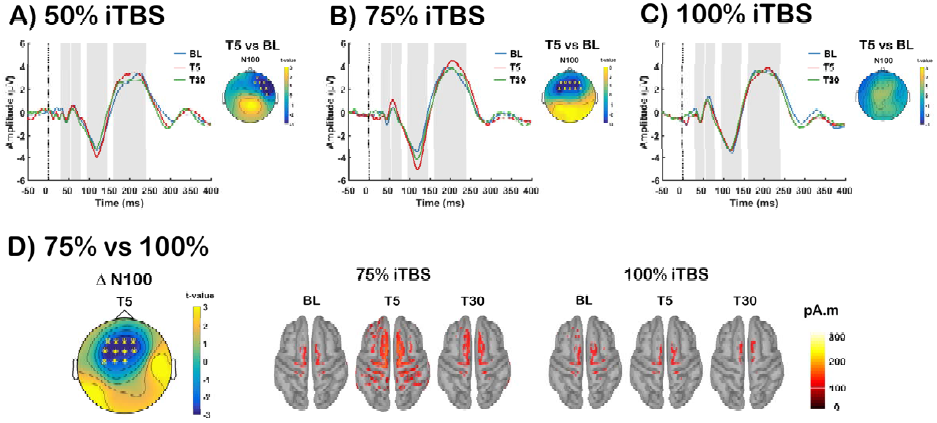
Assessment of transcranial magnetic stimulation (TMS)-evoked potentials (TEPs) before and after each stimulation condition [A: Intermittent theta-burst stimulation (iTBS) at 50% rMT (50% iTBS); B: iTBS at 75% rMT (75% iTBS); C: iTBS at 100% rMT (100% iTBS)]. Grand average TEP waveforms before (BL: blue), 5-min post (T5: red) and 30-min post (T30: green) iTBS at FCz electrode for each stimulation conditions, with significant differences across the scalp illustrated in topoplots. (D) Global scalp differences of iTBS-induced change in N100 amplitude (TEP Δ N100) between 75% and 100% iTBS at T5 and Minimum Norm Estimates (MNEs) of the source level activity at the cortex for the N100 peak. Asterisks and ‘X’s on topoplots indicate significant clusters between comparisons (cluster-based statistics, ^*^*p* < 0.01, ^×^*p* < 0.025).

The differences were not apparent when these conditions were compared with 50% iTBS (all *p* > 0.025), placing the strength of the after-effect of 50% iTBS in the middle of 75% and 100% iTBS.

Given the implication of P60 and N100 peaks in excitatory-inhibitory balance, we explored the relationship between these peaks modulated by iTBS. Correlation analysis was conducted on the data combined across different conditions (n = 48) using the average of 3 fronto-central electrodes (FC1, FCz, FC2) as these electrodes were close to the stimulation, and often showed significant changes following iTBS. Spearman’s rank correlation revealed a significant correlation between Δ P60 and Δ N100 at T5 (r = −0.385, *p* = 0.007) and a trend toward significance at T30 (r = −0.257, *p* = 0.077) (Fig 4).

**Figure 4.**
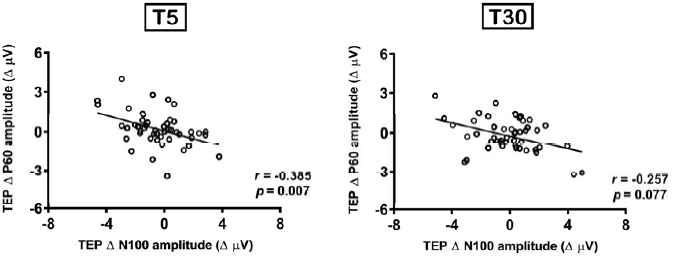
Correlation between intermittent theta burst stimulation (iTBS)-induced changes in transcranial magnetic stimulation (TMS)-evoked potential (TEP) N100 and P60 amplitude

iTBS-induced changes in TMS-evoked oscillations (averaged across all electrodes) are illustrated in Fig 5. We assessed whether different iTBS conditions altered TMS-evoked theta and gamma power (total activity: evoked + induced) in a similar fashion. The cluster-based permutation test revealed a significant increase in TMS-evoked theta power at T5 compared to BL in close proximity to the stimulation site following 75% iTBS (*p* = 0.024), but not with 50% or 100% iTBS (*p* > 0.025). Between conditions, the change in theta power (Δ theta) was larger following 75% iTBS compared to 100% iTBS (75% > 100% iTBS: T5, *p* = 0.020; Fig 5D, top row). However, no prolonged theta change was observed at T30 (all *p* > 0.025).

**Figure 5.**
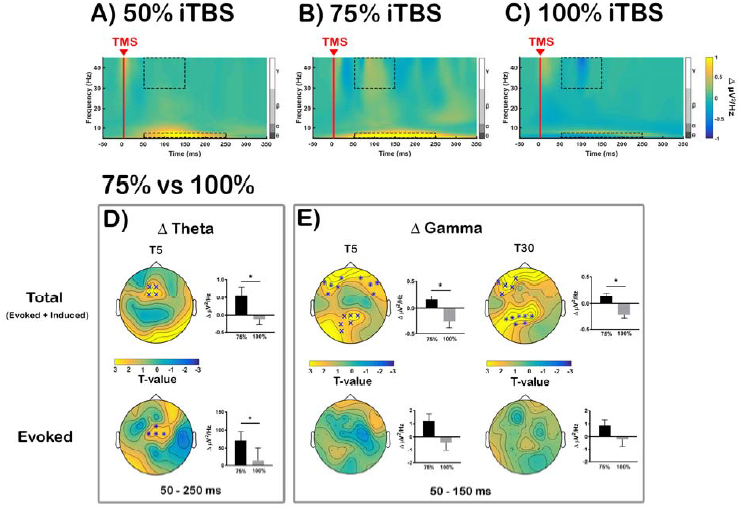
Comparison of transcranial magnetic stimulation (TMS)-evoked oscillations in iTBS-induced changes [A: Intermittent theta-burst stimulation (iTBS) at 50% rMT (50% iTBS); B: iTBS at 75% rMT (75% iTBS); C: iTBS at 100% rMT (100% iTBS)]. Grand average time-frequency plots are illustrated using average of all electrodes. Dotted boxes represent time-frequency windows for gamma (50 – 150 ms) and theta (50 – 250 ms) bands where statistical analyses were conducted. Comparison between 75% and 100% iTBS conditions in (D) Δ theta at T5 and (E) Δ gamma at T5 and T30 across the scalp. Both total power (evoked + induced; top row) and evoked power alone (bottom row) were examined separately. Asterisks and‘X’s on topoplots indicate significant clusters between comparisons (cluster-based statistics, ^*^*p* < 0.01, ^×^*p* < 0.025). Bar graphs were plotted using the values extracted from the significant sensors (when not significant, using same sensors as total power) to examine the directional changes.

Initially, TMS-evoked gamma power showed slightly different changes, with significantly decreased gamma power following 100% iTBS at T5 (*p* = 0.023), which was most pronounced over the frontal sensors. On the other hand, 75% iTBS exhibited non-significant increase in the frontal and parietal regions. Even though no significant differences between BL and T30 were observed in gamma frequency band in any stimulation conditions (all *p* > 0.025), between condition comparisons revealed the change in gamma power (Δ gamma) was significantly different between 75% and 100% iTBS at both T5 and T30, which resulted from polarity-specific changes following the two stimulation conditions. At T5, the difference was most pronounced over bilateral frontal sensors (*p* = 0.006) and parieto-occipital sensors (*p* = 0.015), and at T30, the differences were observed at left frontal (*p* = 0.019) and left parietal region (*p* = 0.006) (Fig 5E, top row). Again, no significant differences were found between 50% and 75% iTBS, or 50% and 100% iTBS for changes in theta or gamma power (all *p* > 0.025).

Examination of evoked oscillations revealed that only 75% iTBS significantly increased both theta (*p* = 0.019, fronto-central) and gamma power (*p* = 0.016, parieto-occipital) at T5, but not at T30. Neither 50% nor 100% iTBS showed any significant change in these frequency bands (all *p* > 0.025). For between condition comparisons, Δ theta showed significant difference between 75% and 100% iTBS at T5 (75% > 100% iTBS: *p* = 0.009, fronto-central) (Fig 5D, bottom row), but not in Δ gamma (Fig 5E, bottom row). No significant differences in Δ theta or Δ gamma were found between 50% and 75% iTBS, or 50% and 100% iTBS at any time point (all *p* > 0.025).

Exploratory analyses including all dimensions of the data (time x frequency x space) were conducted to investigate iTBS-induced changes in all oscillatory bands and time windows. However, we found no significant differences within or between stimulation conditions (all *p* > 0.025). Subgroup analysis on alpha (8 – 12 Hz) and beta (13 – 29 Hz) frequency bands [time x alpha/beta (frequency range averaged prior to cluster-statistics) x space] also resulted in no significant changes (all *p* > 0.025).

### 3.3. The effect of different iTBS intensities on working memory neurophysiology

Before we examined the effect of iTBS on ERPs during WM task, the effect of memory load on the ERPs was first established in our dataset using BL measurement (Supplementary Fig 1A). To test if iTBS-induced changes measured by TEPs were consistent with electrophysiology recordings during WM task, we investigated the ERPs during 2-back and 3-back tasks in a similar manner to TEPs. Supporting the outcome in the TEPs measurement, 75% iTBS significantly increased the amplitude ERP N200 (198 – 218 ms, *p* = 0.022, fronto-central) during 2-back task (Fig 6A). The change in N200 amplitude (ERP Δ N200) was the largest with 75% iTBS compared to 100% iTBS (ERP Δ N200: 190 – 228 ms, *p* = 0.018, fronto-central). Source estimation of ERP N200 in these stimulation conditions revealed activity of parieto-occipital origin, and 75% iTBS resulted in increased activity including fronto-central region, whereas minimal change was observed following 100% iTBS (Fig 6C). Similar to TEPs, the differences were not significant when these conditions were compared with 50% iTBS (all *p* > 0.025). During the 3-back task, cluster-based statistics revealed 50% and 75% iTBS, but not 100% iTBS, resulted in significant differences between BL and T15 in ERP P300 amplitude, which was observed over anterior (50% iTBS: 310 – 333 ms, *p* = 0.023; 75% iTBS: 315 – 343 ms, *p* = 0.015) sensors, indicating increased amplitude following 50% and 75% iTBS (Fig 6B). However, no significant differences were seen between different stimulation conditions in P300 or any other peaks (all *p* > 0.025).

**Figure 6.**
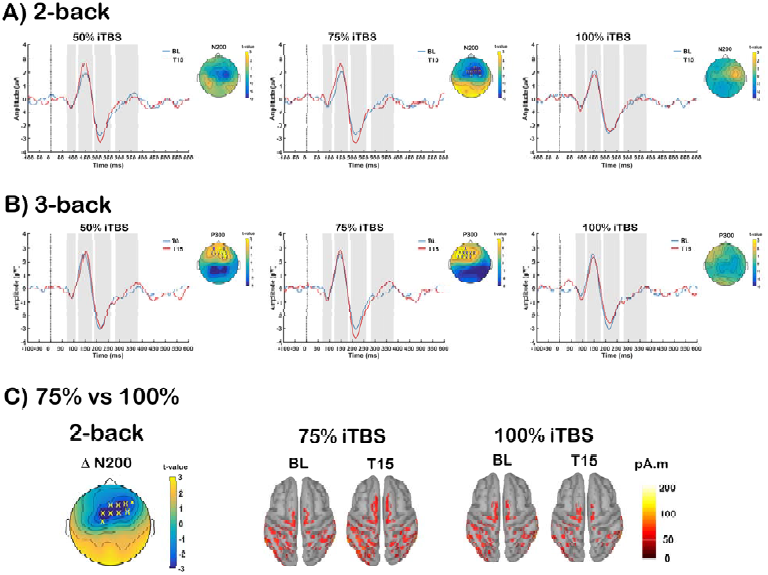
Effect of different intensities of intermittent theta-burst stimulation (iTBS) on the event related potentials (ERPs) during working memory tasks. Grand average ERP waveforms at baseline (BL: blue) and 15-min post (T15: red) iTBS at FCz electrode for each stimulation conditions (50%, 75% and 100% iTBS) in (A) 2-back and (B) 3-back tasks, with significant differences across scalp shown in topoplots. (C) Global scalp differences of iTBS-induced change in N200 amplitude (ERP Δ N200) during 2-back task between 75% and 100% iTBS at T15 and Minimum Norm Estimates (MNEs) of the source level activity at the cortex for the N200 peak. ‘X’s on topoplots indicate significant clusters between comparisons (cluster-based statistics, ^×^*p* < 0.025).

As both the TEP N100 and cognitive task related N200 peaks have been associated with inhibitory mechanisms [TEP N100 (Farzan et al., 2013; Premoli et al., 2014; Rogasch et al., 2015); ERP N200 (Aron, 2007; Kopp et al., 1996; Sasaki et al., 1989)], correlation analysis was performed on the data combined across different conditions (n = 45) using the average of 3 fronto-central electrodes (FC1, FCz, FC2). These electrodes were close to the stimulation, and the significant changes were most often observed in these electrodes across different measures. Spearman’s rank correlation revealed TEP Δ N100 amplitude following iTBS (T5) correlated with ERP Δ N200 amplitude during 2-back task (T15) (r = 0.572, *p* = 0.001; Fig 7). We also explored if TEP Δ N100 correlated with ERP Δ P150 during 2-back task, or ERP Δ 300 during 3-back task, however, no significant correlations were found (all *p* > 0.05). The correlation between TEP Δ N100 and ERP Δ N200 during 2-back task supports the evidence that iTBS alters cortical inhibition in human prefrontal cortex at subthreshold intensities.

**Figure 7.**
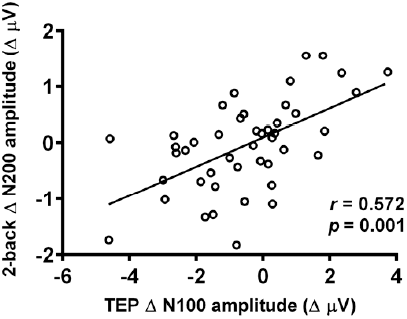
Correlation between iTBS-induced changes in TMS-evoked potential (TEP) N100 amplitude and 2-back task related N200 amplitude

We also assessed the effect of different iTBS intensities on theta and gamma oscillations during WM. The effect of memory load on these oscillations was again examined using BL measurement (Supplementary Fig 1B). As N-back task involves continuous mix of encoding, updating and maintaining of the letters, we divided each trial into two blocks – during letter presentation (50 – 450 ms: encoding) and after letter presentation (550 – 950 ms: maintenance).

To assess whether iTBS was able to modulate these frequency bands during WM task, both theta and gamma power were compared across time, and the changes between conditions. During letter presentation, iTBS did not change any frequency band during 2-back task (all *p* > 0.025). However, during 3-back task, significant increases in theta power were found at T15 compared to BL following both 50% (*p* = 0.013, right prefrontal) and 75% iTBS (*p* = 0.023, left prefrontal), but not 100% iTBS, indicating theta oscillations increased with subthreshold intensities. When Δ theta power were compared between conditions, differences were observed only between 75% and 100% iTBS (75% > 100% iTBS, *p* = 0.022) over left prefrontal sensors (Fig 8A, top row). While gamma power changes were not observed in any stimulation conditions in any N-back task, Δ gamma was significantly different between 75% and 100% iTBS (75% > 100% iTBS, *p* = 0.022) over left posterior sensors during 2-back task (Fig 8B, top row), but not during 3-back task. These findings suggest that iTBS differentially modulates cortical oscillations during letter presentation across task loads. After the letter presentation, however, iTBS resulted in no change in either theta or gamma band during either memory task (all *p* > 0.025), which suggests iTBS was not able to alter the processing involved in maintenance of memory.

**Figure 8.**
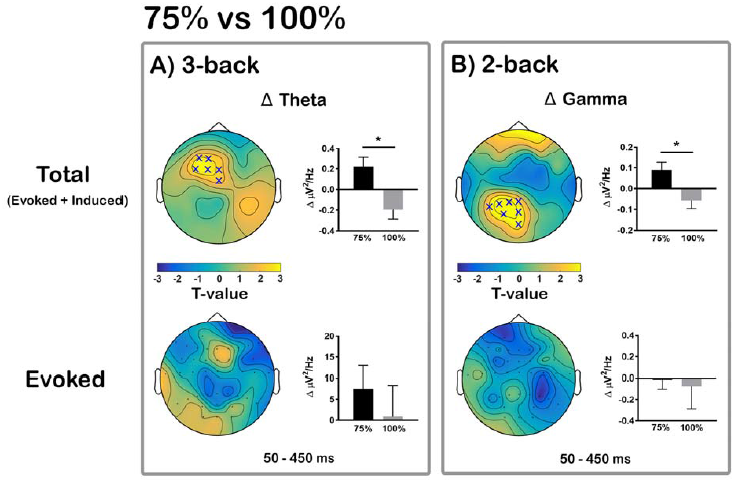
Comparison of intermittent theta-burst stimulation (iTBS)-induced changes in theta and gamma oscillations during different working memory tasks between 75% and 100% iTBS conditions. Significant differences in iTBS-induced change in (A) Δ theta power during 3-back task and in (B) Δ gamma power during 2-back task across the scalp. Both total power (evoked + induced; top row) and evoked power alone (bottom row) were examined separately. ‘X’s on topoplots indicate significant clusters between comparisons (cluster-based statistics, ^×^*p* < 0.025). Bar graphs were plotted using the values extracted from the significant sensors (when not significant, using same sensors as total power) to examine the directional changes.

Analysis of evoked oscillations resulted in a different pattern to the evoked oscillatory activity during TMS-EEG. We found no significant differences in any frequency bands within or between conditions in any task (all *p* > 0.025). However, we observed non-significant increase in theta power following 75% iTBS, which was absent following 100% iTBS in 3-back task (Fig 8A, bottom row). We were unable to detect any changes in gamma power in the evoked activity in 2-back task (Fig 8B, bottom row).

Similar to TMS-EEG time-frequency analyses, exploratory analyses including all dimensions of the data (time x frequency x space) were conducted to investigate iTBS-induced changes in all oscillatory bands and time windows. However, no significant differences were found within or between stimulation conditions both in 2-back and 3-back task (all *p* > 0.025). In addition, we found no significant differences in alpha or beta frequency band (all *p* > 0.025).

We tested if TMS-evoked oscillations (Δ theta and Δ gamma) shared similar mechanisms to N-back task related oscillations (Δ theta with 3-back, Δ gamma with 2-back). For gamma oscillations, average of 3 left/mid parietal electrodes (P3, P1, Pz) were used for correlation analysis as these were found significant in cluster-based analysis of both TMS-evoked and 2-back task. Spearman’s rank correlation revealed the TMS-evoked Δ gamma power following iTBS (T5) correlated with Δ gamma power during 2-back task (T15) (r = 0.420, *p* = 0.004; Fig 9). For theta oscillations, average of 3 fronto-central electrodes (FC1, FCz, FC2) were used. However, no significant correlation was found in Δ theta power between TMS-evoked and 3-back task (r = 0.078, *p* = 0.609).

**Figure 9.**
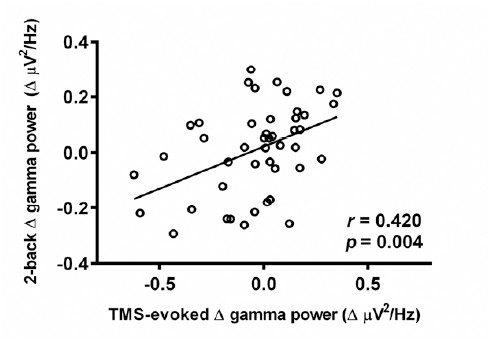
Correlation between iTBS-induced changes in TMS-evoked gamma power and 2-back task related gamma power

### 3.4. The effect of iTBS intensity on working memory

N-back WM performance (accuracy d’, reaction time and effect sizes (Hedges’ g (Hedges, 1985)) is shown in Table 1.

**Table 1.**
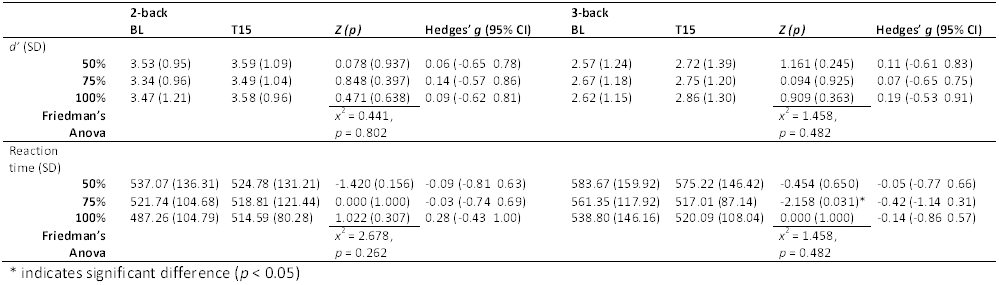
Mean (SD) *d*’, accurate reaction time (ms) and the effect sizes (hedges’ *g*) of 2-back and 3-back after different stimulation conditions.

*Performance at baseline:* Initial statistical analysis was conducted on pre-iTBS (BL) data (combined across sessions, n = 45) to determine if WM performance differed between different memory load conditions (2-back vs 3-back) in accuracy (*d*’) and reaction time. The Wilcoxon signed-rank tests revealed d’ scores decreased [Z = −5.108, *r* = −0.35, *p* = 0.001 (3-back < 2-back)] and reaction times increased [Z = 2.523, r = 0.18, *p* = 0.012 (3-back > 2-back)] with increasing WM load. We also conducted order effect analysis to confirm the effectiveness of the counter-balancing of stimulation conditions. Friedman’s ANOVA showed no significant session effects in either d’ (2-back: x^2^ = 0.037, *p* = 0.982; 3-back: x^2^ = 0.036, *p* = 0.982) or accurate reaction time (2-back: x^2^ = 3.448, *p* = 0.178; 3-back: x^2^ = 1.793, *p* = 0.408) for WM tasks at baseline measure.

*Performance following iTBS*: Following 75% iTBS there was a significant decrease in reaction time (Wilcoxon signed rank test; *p* = 0.031) of small-to-moderate effect size (−0.42) during 3-back task. No other stimulation conditions showed any significant differences in WM performance (Wilcoxon signed rank test; all *p* > 0.05).

*Comparison pre- and post-iTBS:* When compared across conditions using the change-from-baseline scores (post – pre; Δ), we could not detect any significant differences in reaction time or d’ (Friedman’s ANOVA; all *p* > 0.05).

We next tested if physiological changes were related to improved reaction time following 75% iTBS. The correlation analyses were performed between significant changes observed following 75% iTBS in TMS-EEG (Δ N100, Δ theta, Δ gamma) and during 3-back task (Δ P300, Δ theta) against Δ reaction time in 75% iTBS condition during 3-back task. However, there was no significant correlation between any change in physiological measure and 3-back reaction time (all *p* > 0.05).

## 4. Discussion

This study examined the link between iTBS intensity and LTP-like neural plasticity, and the association to neurophysiological and behavioural metrics of WM in the prefrontal cortex. The data indicate an inverse U-shaped relationship between iTBS intensity and neurophysiological changes following single-pulse TMS and during working memory, whereby these effects were maximal at an intermediate intensity of 75% rMT. While the plastic effects correlated with changes in neurophysiological aspects of cognition, they did not translate to behavioural outcomes of WM performance. The data suggest using subthreshold intensities is important in order to achieve desirable neurophysiological after-effects following iTBS in the prefrontal cortex, and highlight potential benefits in the application of iTBS for clinical treatment.

### 4.1. Influence of iTBS intensity on plastic effects in DLPFC

iTBS modulated N100 amplitude, and the increase in this component was maximal when iTBS was delivered at 75% rMT compared to lower (50% rMT) or higher intensity (100% rMT). These findings raise interesting aspects of the relationship between intensity and plasticity induction. Increased N100 following iTBS over prefrontal cortex is in line with previous studies which also showed modulation of this component following iTBS (Chung et al., 2017; Harrington and Hammond-Tooke, 2015) or cTBS (Harrington and Hammond-Tooke, 2015; Huang and Mouraux, 2015), however, opposite outcome (i.e. decreased following iTBS, increase following cTBS) has also been described in cerebellar stimulation (Casula et al., 2016). The N100 deflection is considered to be the most prominent and robust TMS-EEG component and is understood to have the greatest inter-individual and inter-session reproducibility compared to other TEPs both in motor and prefrontal cortex (Lioumis et al., 2009). The N100 is also considered to have a high degree of sensitivity to small changes in cortical excitability (Nikulin et al., 2003). These factors enhance the value of N100 as a marker of cortical processing in basic and clinical research, and make it ideal for exploration of the effects of TMS plasticity paradigms (Chung et al., 2015b; Ilmoniemi and Kicic, 2010; Noda et al., 2016). Recent studies have provided evidence that N100 may also be associated with GABA_B_-mediated postsynaptic inhibition in motor (Farzan et al., 2013; Premoli et al., 2014; Rogasch et al., 2013a) and prefrontal (Rogasch et al., 2015) cortex. These findings raise the prospect of current data reflecting an increase in cortical inhibition following iTBS. This account is difficult to reconcile with the absence of effects of iTBS on GABA_B_-mediated inhibitory measures such as long intracortical inhibition (LICI) (Goldsworthy et al., 2013; Suppa et al., 2008) or the cortical silent period (Brownjohn et al., 2014; Di Lazzaro et al., 2011) in the motor cortex. However, we did recently observe an increase in LICI of theta oscillations which correlated with increased N100 amplitude following iTBS over DLPFC (Chung et al., 2017), supporting possible modulation of cortical inhibition following stimulation.

The present data did not show a significant increase in P60 amplitude following iTBS. Recent studies suggest that P60 provides a marker of neural excitability (Cash et al., 2017b; Hill et al., 2017). It should be noted that the SNR for P60 is substantially lower than for N100, and the current protocol with ∼ 47 single TMS pulses may have been insufficient to capture significant changes. The data, however, demonstrated evidence of a relationship between the change in amplitude of N100 and P60 following iTBS. If N100 is related to inhibition, and P60 to neural excitability, this finding suggests that the change in excitation was balanced by a similar change in inhibition, maintaining the excitatory-inhibitory balance following iTBS. This is in agreement with the concept of homeostatic plasticity mechanisms involving a dynamic adjustment of excitatory and inhibitory circuits (Turrigiano and Nelson, 2004). In summary, it appears that the most reliable TMS-EEG metric of plasticity is the modulation of N100 amplitude and iTBS-induced change in this component was greatest at an intermediate intensity of 75% rMT.

The relationship between iTBS intensity and the level of plasticity induction is likely explained by the unique mechanistic features underlying iTBS. Typically, the propensity for LTP-like effects increases with increasing intensity (Artola et al., 1990; Cash et al., 2017a), whereby greater postsynaptic depolarisation leads to higher levels of N-methyl-D-aspartate receptor (NDMA-R) activation, and consequently regulating the processes leading to LTP (Luscher and Malenka, 2012). A similar relationship has been demonstrated across a range of non-invasive NMDA-R dependent brain stimulation protocols in human (Batsikadze et al., 2013; Cash et al., 2017a; Doeltgen and Ridding, 2011; Moliadze et al., 2012). However, the present findings demonstrate an exception to this relationship, showing an inverse U-shaped influence of stimulus intensity on plastic effects. This may be explained by the unique temporal aspects that underlie the fundamental mechanism of TBS (Larson and Munkacsy, 2015). It is thought that the robust after-effect of TBS is achieved through targeting a late period of presynaptic GABA_B_-mediated disinhibition, which may itself help sustain the theta rhythm (Davies et al., 1991; Larson and Munkacsy, 2015; Mott and Lewis, 1991). More specifically, stimulation elicits both postsynaptic GABA_B_-mediated inhibition (inhibitory postsynaptic potentials) and presynaptic GABA_B_ autoreceptor-mediated disinhibition (temporary blockade of further GABA release). It has been shown that presynaptic disinhibition outlasts postsynaptic inhibition, resulting in a late temporal window (∼ 200 ms) during which disinhibition dominates (Deisz, 1999; Otis et al., 1993) and plasticity induction is enhanced (Davies and Collingridge, 1996; Larson and Lynch, 1986; Mott and Lewis, 1991; Pacelli et al., 1989). Delivery of stimulus bursts at this interval (i.e. TBS) results in a rapid induction of plastic effects (Davies et al., 1991; Mott and Lewis, 1991). A similar late phase of disinhibition has recently been described in humans at ∼ 200 ms latency (Cash et al., 2010), during which excitability (Cash et al., 2011) and plasticity induction were enhanced (Cash et al., 2016). Importantly, the latency of this period increases with increasing stimulus intensity (Cash et al., 2010) and stimulation outside this window does not result in plastic effects in humans (Cash et al., 2016) or animals (Larson and Munkacsy, 2015). Consequently, higher TBS intensities may miss this plastic window. This unique plasticity mechanism may account for the inverse U-shaped relationship between intensity and plasticity observed in this study.

### 4.2. The effect of iTBS on neural oscillations is modulated by stimulus intensity

The spectral characteristics elicited by single-pulse TMS are commonly modulated following TBS in a manner that may depend on the area being stimulated. Cerebellar stimulation (iTBS and cTBS) were found to modulate alpha and beta power (Casula et al., 2016), while cTBS of motor cortex produced modulation of theta, alpha and beta power (Vernet et al., 2013). In the prefrontal cortex, polarity-specific changes in TMS-evoked theta power (increase following iTBS, decrease following cTBS) were demonstrated (Chung et al., 2017), and modulation of theta and gamma power were also observed in a resting EEG study (Wozniak-Kwasniewska et al., 2014), suggesting TBS may be targeting the natural frequency of oscillations in the stimulated region. Our data indicate the additional dimension of iTBS intensity in modulating these spectral changes. Theta power was increased following iTBS, consistent with our previous study (Chung et al., 2017), and this effect was maximal at 75% rMT. Previous findings in relation to gamma power have been somewhat inconsistent, showing no change (Chung et al., 2017), or an increase following cTBS (Vernet et al., 2013), and this may relate to low SNR of gamma and/or discrepancies in analysis methods such as the total power vs evoked power, and the level of spatial dynamics (region of interest vs global scalp analysis). Here, we examined both total and evoked activity, and the analysis of total power provided additional information about the spread of activity following iTBS in distant yet interconnected regions. In the present study, the direction of change in TMS-evoked gamma was further shown to depend on the intensity of the stimulation (increase with 75% iTBS, decrease with 100%), and this change remained significant at T30. This was an interesting observation as iTBS on rat cortex also resulted in long-lasting gamma power increase (Benali et al., 2011), and this finding may indicate that the persistent difference in after-effects of iTBS could be observed in the gamma frequency band in humans. The increase in theta and gamma power following iTBS at 75% rMT would seemingly suggest that this intensity might be advantageous for enhancing performance on cognitive tasks. We did not find this, however.

### 4.3. Relationship to neurophysiological metrics during cognitive performance

Similar to TMS-EEG findings, iTBS modulated neurophysiological metrics during the performance of the cognitive task in an intensity-dependent manner. With 75% rMT and 50% rMT to some extent, iTBS increased N200 amplitude in the 2-back task, and P300 amplitude in the 3-back task, while no changes were evident at a higher intensity. N200 has been linked to executive control (Kopp et al., 1996) and cognitive and inhibitory processing (Folstein and Van Petten, 2008; Sasaki et al., 1989; Schmajuk et al., 2006). The change in ERP N200 amplitude correlated with plastic changes in TEP N100 amplitude, suggesting that these may be modulated by iTBS in a similar manner or have a degree of functional overlap. This link was not present with ERP P150 or ERP P300, further strengthening the selective link for possible inhibitory processing involved in two different measures following iTBS. In the frequency domain, frontal theta power was enhanced following 75% iTBS during the 3-back task. There was also a trend for an increase in parietal gamma power during 2-back WM task, which was maximal following iTBS at 75% rMT. These results are consistent with a maximal effect of iTBS at 75% rMT observed in TMS-EEG data. A significant correlation between the change in TMS-evoked gamma power and event-related gamma power during 2-back task provides further evidence of a relationship between the neural elements modulated by TBS, probed by single-pulse TMS and functionally recruited during a WM task. These results support the notion that iTBS Mcan likely enhance the neurophysiological mechanisms mediating working memory (Hoy et al., 2015), and does so in an intensity-dependent manner. Theta and gamma oscillations are important in WM (Howard et al., 2003; Hsieh and Ranganath, 2014) and these oscillation frequencies are targeted by iTBS. The involvement of fronto-parietal network control system (Dosenbach et al., 2008) is also supported by the observation of the influence of TBS on visuospatial attention (Xu et al., 2013) and in WM task (Hoy et al., 2015). Theoretically, one would expect these physiological changes to have an impact on behavioural outcome.

### 4.4. Marginal influence of iTBS on working memory performance

Despite robust changes in TMS-evoked potentials, and WM-related ERPs and theta oscillations, no significant differences were observed in WM performance between stimulation conditions. This is contrary to a previous finding which demonstrated a significant increase in the accuracy of 2-back task following iTBS compared to sham stimulation (Hoy et al., 2015). The reason for this discrepancy remains unclear, and further research is required as currently only a few studies have been performed in this area to date (Cheng et al., 2016; Debarnot et al., 2015; Demeter et al., 2016; Hoy et al., 2015; Ryals et al., 2016). However, a similar lack of behavioural changes in the presence of robust neurophysiological effects after tDCS has also been described with a larger sample size (n = 20) (Hill et al., 2016), suggesting neurophysiological measures may provide a more sensitive index for assessing changes following neuromodulatory paradigms. This points to a possible conclusion that the inadequate behavioural findings in this study may be due to a ceiling effect of performance in healthy individuals. As such, the ability for iTBS to enhance accuracy may have been limited, and thus a significant level of behavioural change was unattainable. A recent meta-analysis of the working memory performance following non-invasive brain stimulation demonstrates only small effect sizes in improvement in healthy controls compared to clinical populations that showed medium effect sizes (Brunoni and Vanderhasselt, 2014). Greater behavioural effects may be detected in disorders of WM, such as schizophrenia, in which considerable differences in physiological measures are often observed compared to a control group (Ferrarelli et al., 2012; Noda et al., 2017). Therefore, the lack of behavioural outcome in this study is potentially consistent with neurophysiological findings, where we only observed decreased reaction time following iTBS at 75% rMT.

### 4.5. Limitations

Our study design did not include a sham condition, however, this was because the primary intention was to investigate intensity-dependent effects of TBS. Nonetheless, the effects of 75% iTBS in this study closely resemble those of our previous sham-controlled study (Chung et al., 2017). The lack of performance changes in our healthy subjects may reflect ceiling effects and it is possible that clinical cohorts may show greater cognitive benefits. Our TEP data mainly indicate changes in the TEP components which had a high SNR (Supplementary Table S1). It is possible that increasing the number of stimuli would also have revealed changes in other TEP components. Finally, consistency of stimulation site localisation between sessions could be improved by using MRI-guided neuronavigation, however, this was not feasible in the present study. Nevertheless, the TEP waveforms in this study are consistent with previous studies in DLPFC (Chung et al., 2017; Hill et al., 2017; Rogasch et al., 2014), and comparable results have been reported using EEG-guided methods (Rogasch et al., 2014) and MRI-guided neuronavigation (Lioumis et al., 2009).

## 5. Conclusion

The present data provide the first evidence that for iTBS, unlike rTMS (Nahas et al., 2001; Padberg et al., 2002), using higher intensities may not be optimal for maximal neuromodulation, and instead, maximal effects are observed at an intermediate intensity of 75% rMT. Further research is required to explore whether the present intensity relationship extends to clinical efficacy. The data also indicate that the link between neurophysiological and behavioural effects may not be as direct as hoped, however, it is also possible that repeated sessions are necessary to elicit more robust behavioural outcomes. Despite the modest behavioural outcome in this study, the change in cortical oscillatory activity evoked by TMS has been implicated in clinical improvement in a patient with MDD (a case study) (Pellicciari et al., 2017a), and we were able to demonstrate different effects the stimulation intensity has on the oscillatory properties following iTBS over the left prefrontal cortex, which may be useful indices for treatment regime. Other short paradigms of similar duration to TBS are now available (Cash et al., 2017a; Cash et al., 2016), and may offer another option for clinical trials.

In conclusion, the current study indicates that iTBS at 75% rTMS produces the strongest effect on physiological measures in the prefrontal cortex, and increasing the intensity may not necessarily result in a corresponding change. These findings highlight the importance of intensity in administering iTBS and paves the path for more efficacious outcome in patients with neurological and psychiatric disorders.

## Supplementary results

### SNR analysis

Table S1 shows the SNR of single-pulse TMS before and after different iTBS conditions. Mean SNR values were averaged across individual for grand average values. Values greater than 3 SDs (99.7% of the baseline distribution) are considered as good SNR. Qualitatively only N45 peaks showed moderate values (∼2.5 SDs), but other peaks, especially latter peaks (N100 and P200) showed excellent SNR.

## Neurophysiology of different working memory load

### Event-related potentials (ERPs)

In order to establish the effect of memory load on the ERPs in our dataset, each task performed before iTBS (BL) was combined across sessions (n = 45) and the amplitude of ERPs during 2-back and 3-back tasks were compared. Visual representations during 2-back and 3-back tasks resulted in a series of consistent negative [N100 (∼90 ms) – associated with discrimination processing (Itier and Taylor, 2004); N200 (∼215 ms) – attention and inhibition (Coull, 1998; Kopp et al., 1996)] and positive peaks [P150 (∼145 ms) – associated with perceptual priming mechanism (Gosling et al., 2016); P300 (∼350 ms) – availability of processing resources (McEvoy et al., 1998)] at FCz electrode (Fig 4A). Cluster-based statistics across space revealed significant differences around these peaks [N100 (3-back < 2-back) over left fronto-temporal (*p* = 0.002) and right posterior sensors (*p* = 0.002); P150 (3-back > 2-back) over anterior (*p* = 0.0004) and posterior sensors (*p* = 0.002); N200 (3-back > 2-back) over anterior (*p* = 0.002) and posterior sensors (*p* = 0.016); P300 (3-back < 2-back) over fronto-central sensors (*p* = 0.003)] (Supplementary Fig 1A).

### Event-related oscillations

To test if the power of these frequencies differed based on memory load, BL measures were again combined across sessions (n = 45) and the power of theta and gamma bands was compared between 2-back and 3-back task. As N-back task involves continuous mix of encoding, updating and maintaining of the letters, we divided each trial into two blocks – during letter presentation (50 – 450 ms: encoding) and after letter presentation (550 – 950 ms: maintenance).

During letter presentation, cluster-based permutation tests revealed an increase in theta power over left frontal sensors (*p* = 0.024), and in gamma power over frontal (*p* = 0.0004) and posterior sensors (*p* = 0.0004) in 3-back compared to 2-back conditions (Fig. 6A). After letter presentation, more prominent increases were observed in theta power (*p* = 0.022, left frontal; *p* = 0.002, posterior) in 3-back conditions, whereas less pronounced increases were also observed in gamma power (*p* = 0.030, right frontal; *p* = 0.038, left posterior) (Supplementary Fig 1B).

---

**Table S1.**
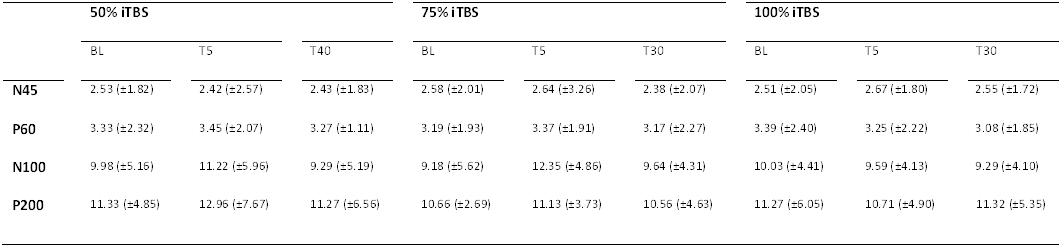
Signal-to-noise ratio (SNR) of each peak before (BL), 5-min post (T5) and 30-min post (T30) stimulation (mean ± SD)

**Figure S1.**
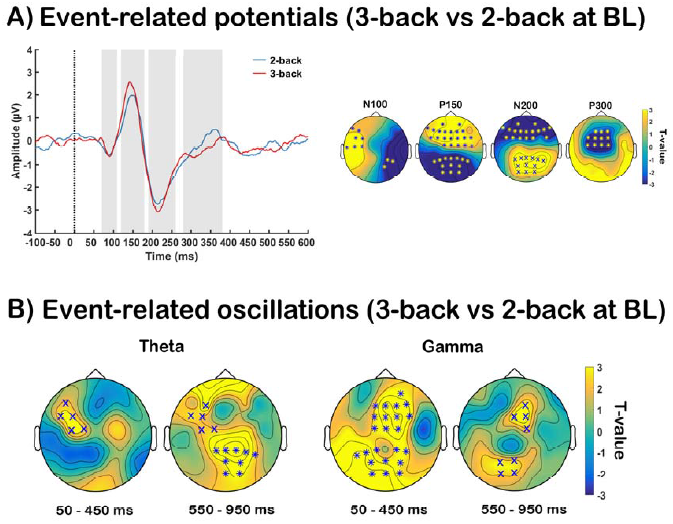
Comparison of event-related potentials and oscillations between different memory loads during working memory tasks at baseline (BL). (A) Grand average ERP waveforms at FCz electrode and (B) differences in theta and gamma power between 2-back (blue) and 3-back (red) tasks before iTBS, with significant differences across the scalp illustrated in topoplots (yellow / blue: 3-back more positive / negative than 2-back). Asterisks and ‘X’s on topoplots indicate significant clusters between comparisons (cluster-based statistics, \**p* < 0.01, ^×^*p* < 0.05).

